# Immunomodulatory Extracellular Matrix Hydrogel Mitigates Scar Formation in a Model of Tongue Fibrosis

**DOI:** 10.1101/2022.05.17.492250

**Authors:** EI Zelus, A Panduro, M Alperin, AM Vahabzadeh-Hagh, KL Christman

## Abstract

**Background:** While head and neck cancer treatment regimens, including surgical resection, irradiation, and chemotherapy, are effective at removing tumors, they lead to muscle atrophy, denervation, and fibrosis, contributing to the pathogenesis of tongue dysphagia – difficulty swallowing. Current standard of care is ineffective; we propose an alternative approach utilizing an acellular and minimally invasive biomaterial to preserve muscle content and reduce fibrosis of the tongue after injury. Here, we investigate a decellularized extracellular matrix hydrogel for the treatment of tongue fibrosis in a partial glossectomy injury model.

**Methods:** Skeletal muscle extracellular matrix (SKM) hydrogel was fabricated by decellularizing porcine skeletal muscle tissue through established protocols. A partial glossectomy injury in the rat was used as a model of tongue fibrois, and SKM hydrogels along with saline controls were injected to the site of scarring two weeks after injury. Tissues were harvested at 3 and 7 days post-injection for gene expression analysis of immune and myogenic pathways, and at 4 weeks post-injection to evaluate histomorphological changes in skeletal muscle and scar formation.

**Results:** SKM hydrogel reduced scar formation and improved muscle fiber cross-sectional area in the region of injury compared to saline controls. SKM upregulated pro-regenerative immune response while downregulating pro-inflammatory response and further promoted angiogenic gene expression.

**Conclusion:** This study demonstrates the immunomodulatory and tissue-regenerative capacity of an acellular and minimally invasive biomaterial in a rodent model of tongue fibrosis.

## Introduction

Dysphagia is a debilitating condition defined broadly as difficulty swallowing. Dysphagia may result from atrophy, denervation, or fibrosis of the tongue muscles.^1^ While this condition is commonly implicated in aging or neurological disorders, another patient population of interest is those recovering from head and neck cancer.^2^ Treatment of these cancers often involves surgical removal of the tumor followed by radiation with or without chemotherapy. About 54,000 cases of oropharyngeal cancer arise annually in the US, with approximately 11,000 deaths.^3^ Dysphagia is a common sequela of treatment, affecting at least half of head and neck cancer patients.^4^ Dysphagia is a morbid condition that severely affects patient quality of life, and can lead to feeding tube dependence, aspiration, malnutrition, and even death.^5,6^ Current standard of care for oropharyngeal dysphagia is limited to rehabilitative strategies such as swallow therapy and lingual muscle exercises. However, these approaches do not provide long-term improvement in swallowing and tongue strength,^7^ and they do not reverse the tissue-level pathology by improving muscle regeneration and reducing scar tissue formation.^7^

Considering these deficiencies in treatment of tongue dysphagia following head and neck cancer treatment, a number of investigative cell-based therapies that seek to regenerate tongue muscle or to provide structural augmentation to improve tongue function have emerged. While preclinical investigation of allogeneic mesenchymal stem cell injection in an athymic rat model has demonstrated some potential,^8,9^ the manufacturing, cost, and difficulties associated with a living stem cell product pose substantial translational challenges. Additionally, autologous muscle-derived cell therapy demonstrated safety but lacked efficacy after 2 years in a Phase 1 trial.^10^ Overall, the current body of research demonstrates a continued need for an accessible and minimally invasive therapeutic that can induce tissue regeneration to reduce scar formation and muscle atrophy to improve muscle repair.

The use of decellularized extracellular matrix (ECM) hydrogels that induce immune modulation, cellular recruitment and differentiation, neovascularization, and ECM remodeling has been explored as a therapeutic in numerous disease phenotypes.^11^ The ECM is a complex network of proteins vital for structural support and cell signaling; when a whole tissue is decellularized, the ECM remains as an acellular biomaterial with tissue regenerative properties.^11^ Additionally, due to their thermoresponsive properties, ECM hydrogels can be delivered minimally invasively via injection, and after exposure to physiologic conditions, the liquid forms a hydrogel comprised of a nanofibrous ECM scaffold.^12^ With sufficient decellularization, the acellular xenogeneic ECM hydrogels are biocompatible and have been utilized in large^13,14^ and small animal models^15^ as well as a human Phase 1 clinical trial that assessed safety and feasibility for intervention in subacute and chronic myocardial infarction.^16^ We have previously demonstrated that a decellularized porcine skeletal muscle ECM hydrogel (SKM) increased vascularization, enhanced the recruitment and differentiation of muscle progenitors, and reduced muscle atrophy and cell death in an ischemic injury model.^17,12,18^ SKM has further been shown to prevent skeletal muscle atrophy and mitigate fibrotic degeneration, as well as modulate the immune response after mechanical muscle injury.^19^

In the present study, we investigated the therapeutic efficacy and potential mechanisms of action of SKM in a preclinical animal model of tongue fibrosis. We observed that SKM reduces scar formation and improves muscle fiber area in a rat tongue partial glossectomy model. We further demonstrated that SKM modulates the immune response and upregulates genes related to angiogenesis. Overall, we found that SKM injection is a promising treatment in a preclinical model of tongue fibrosis that warrants further investigation as a potential minimally invasive acellular therapeutic for oropharyngeal dysphagia.

## Materials and Methods

### SKM Hydrogel Fabrication

SKM material was fabricated as previously described.^20^ Briefly, porcine skeletal muscle was chopped into small cubes before being spun in a 1% sodium dodecyl sulfate detergent solution for 5 days, with daily solution changes. Tissue was thoroughly rinsed in water to remove residual detergent. Decellularized tissue was frozen at −80°C and then lyophilized for 48 hours, after which the material was milled to a fine powder. This powder was partially enzymatically digested with pepsin for 48 hours, and then pH was neutralized, and ionic concentration balanced. SKM was brought to a concentration of 6 mg/mL, which was determined as the optimal concentration for skeletal muscle injection.^21^

### Partial Glossectomy Model

All procedures were approved by the Institutional Animal Care and Use Committee at the University of California, San Diego. Male Sprague Dawley rats weighing between 225-250 g (approximately 3 months old) underwent partial glossectomy injury, as previously established.^9^ Briefly, animals were anesthetized with isoflurane, and buprenorphine was delivered subcutaneously for pain management. The tongue was retracted with a 4–0 silk suture. A 4-mm dermal punch was used to excise a quarter of the tongue, anterior to the circumvallate papillae on the left side. Silver nitrate chemical cautery was used for hemostasis. The animals were then monitored for 5 days postoperatively; the rats were provided with a soft diet for the duration of the study period. Scar formation at the site of injury occurred over the following 2 weeks,^9^ at which time point SKM was injected directly into the tongue.

### Reliability of the SKM Injection and Volume Optimization

To enable visualization of SKM *in situ*, SKM was prelabeled with Alexa Fluor™ 568 NHS Ester (Thermo Fisher Scientific, Waltham, MA). SKM was incubated with the dye on ice for an hour to ensure complete binding. Fluorescently pre-labelled SKM was injected directly into the site of injury, two weeks following a partial glossectomy injury. Injection volumes of 50, 100, 200, and 300 μL were tested (n=2 animals per volume); this range was determined based on SKM injections in other skeletal muscle injury models^21^ and cell injections in the rat tongue^9^. Animals were euthanized and tongues were harvested 1 week following injection; cryosectioned and stained with DAPI nuclear counterstain for tissue localization. Tissue cross-sections were then visualized at 20X magnification using a Leica Ariol® fluorescent microscope.

### Histomorphological Assessment of SKM Therapeutic Efficacy

Two weeks following partial glossectomy injury, SKM or saline were injected into the injury site. Based on the injection volume optimization study, the 200 and 300 μL demonstrated good retention and spread in the tongue tissue. Thus, experimental groups included 200 μL SKM, 300 μL SKM, and 200 μL saline (n=6/group). Animals were euthanized 4 weeks following injection and tongues were harvested and cryosectioned. Masson’s Trichrome (Polysciences, Warrington, PA) stain was used to identify collagen and muscle, and tissue sections were visualized using a Leica Aperio ScanScope ® CS2. For quantification of muscle fibers within the scar area, fibrosis was identified with an anti-collagen antibody (Bio-Rad, Hercules, CA, 1:500) with an Alexa Fluor™ 488 secondary (Invitrogen, Carlsbad, California, 1:500) secondary and myofiber membranes were identified with anti-α-sarcoglycan antibody (Leica Biosystems, Wetzlar, Germany, 1:200) with an Alexa Fluor™ 568 secondary (Invitrogen, Carlsbad, California, 1:500). Tissue sections were visualized using a Leica Ariol® fluorescent microscope.

For quantification of the scar area from the Masson’s Trichrome stained tissue sections, Aperio ImageScope software was used to trace the border of the scar region, as defined by the blue collagen stain. The outer edge of the tongue cross-section was traced, and the area of the scar region was normalized to total tongue area for each section. All sections containing the scar were analyzed, and normalized scar area data were averaged per animal. For quantification of muscle fibers from immunohistochemical staining, the border of the scar was identified with an anti-collagen antibody. Then, cross-sectional muscle fibers within the scar region were identified with the anti-α-sarcoglycan antibody and fiber areas were quantified. Muscle fibers with centralized nuclei (identified with DAPI stain) were also identified. Numbers of fibers were normalized to scar area, and the numbers of centrally nucleated fibers were normalized to muscle fiber counts. All sections containing scar were analyzed, and data were averaged per animal.

### RNA Isolation and Nanostring Multiplex Gene Expression Analysis

Two weeks after partial glossectomy injury, either 300 μL of SKM or Saline was injected into the injury site (n=6/group). 300 μL injection volume was chosen based on histological data, and the control saline injection was increased to 300 μL to match SKM. An injured non-injected group was used as an additional control to further evaluate potential effects of the injection itself. At 3 and 7 days post-injection, physiologically relevant timepoints for the immune response and early muscle regeneration, tongues were harvested and submerged in RNAlater™, then stored at 4°C overnight before being transferred to −80°C to preserve tissues for RNA isolation. For RNA isolation, tongue specimens were thawed and tissue was trimmed to isolate the scar region. Isolated scar region was divided in half, to accommodate spin column capacity, and homogenized (TissueRuptorII, Qiagen, Germantown, Maryland), and RNA was isolated with RNAeasy Fibrous Tissue Mini Kit following manufacturer instructions (Qiagen, Germantown, Maryland).

As previously described,^19^ we used a NanoString nCounter ® MAX Analysis System with an nCounter® custom CodeSet of 145 genes involved in pathways relevant to skeletal muscle regeneration.^19^ Briefly, RNA concentration was measured using a Qubit 3.0 Fluorometer with a Qubit™ RNA HS Assay kit. The hybridization buffer (70 μL) was then mixed with the Custom Reporter CodeSet solution, and 8 μL of this master mix was then added to 50-100 ng of RNA per tissue sample, and RNA-free water up to 13 μL total. Then, 2 μL of Capture ProbeSet was added to the mixture, thoroughly mixed and placed on a thermocycler at 65°C for 16-48 hours and then maintained at 4°C for less than 24 hours. Using a two-step magnetic beads purification, probe excess was removed in PrepStation and target/probe complexes were bound on the cartridge. The data were collected by the digital analyzer (NanoString nCounter® Digital Analyzer) with images of immobilized fluorescent reporters in the sample cartridge. Results of barcode reads were analyzed by nSolver™ Analysis Software 4.0, and differential expression analysis was done with a custom R script. The NanoString data were visualized using ggplot and pheatmap packages in R.

### Statistical Analysis

The initial SKM dosing study was a pilot study that did not aim to achieve statistically significant differences, so n=2 animals per injection volume were used for this initial investigation. For the investigation of histomorphological changes following SKM injection, sample size was based on a previous study of SKM for muscle regeneration in a model of hindlimb ischemia;^17^ using G*Power, 6 animals/group were needed to achieve 90% power and a significance of 0.05. For gene expression studies, preliminary data with qRT-PCR were used to calculate the sample size; 11 animals/group/timepoint were necessary to achieve 80% power and a significance of 0.05. Data that followed a parametric distribution were compared using a Student’s t-test, or a one-way analysis of variance followed by Tukey’s post hoc pairwise comparisons. Data that did not follow a parametric distribution (skeletal muscle cross-sectional fiber area) were analyzed by Mann-Whitney test.^19^ Data were analyzed using GraphPad Prism v8.0, San Diego, CA. Gene expression normalization and differential expression was analyzed using the NanoStringDiff package in R, with a significance at a p<0.05.^19^

## Results

### Determining optimal volume of SKM hydrogel

Acellular biomaterial therapeutics have not been previously investigated for oropharyngeal dysphagia in a pre-clinical animal model, so we first conducted a study to determine the optimal injection volume of SKM hydrogel. A rat partial glossectomy injury model, previously developed to mimic the fibrosis and muscle atrophy that follows head and neck cancer treatment, was used.^9^ The study design (Figure 1A) included injection volumes ranging from 50 to 300 μL.

**Figure 1.**
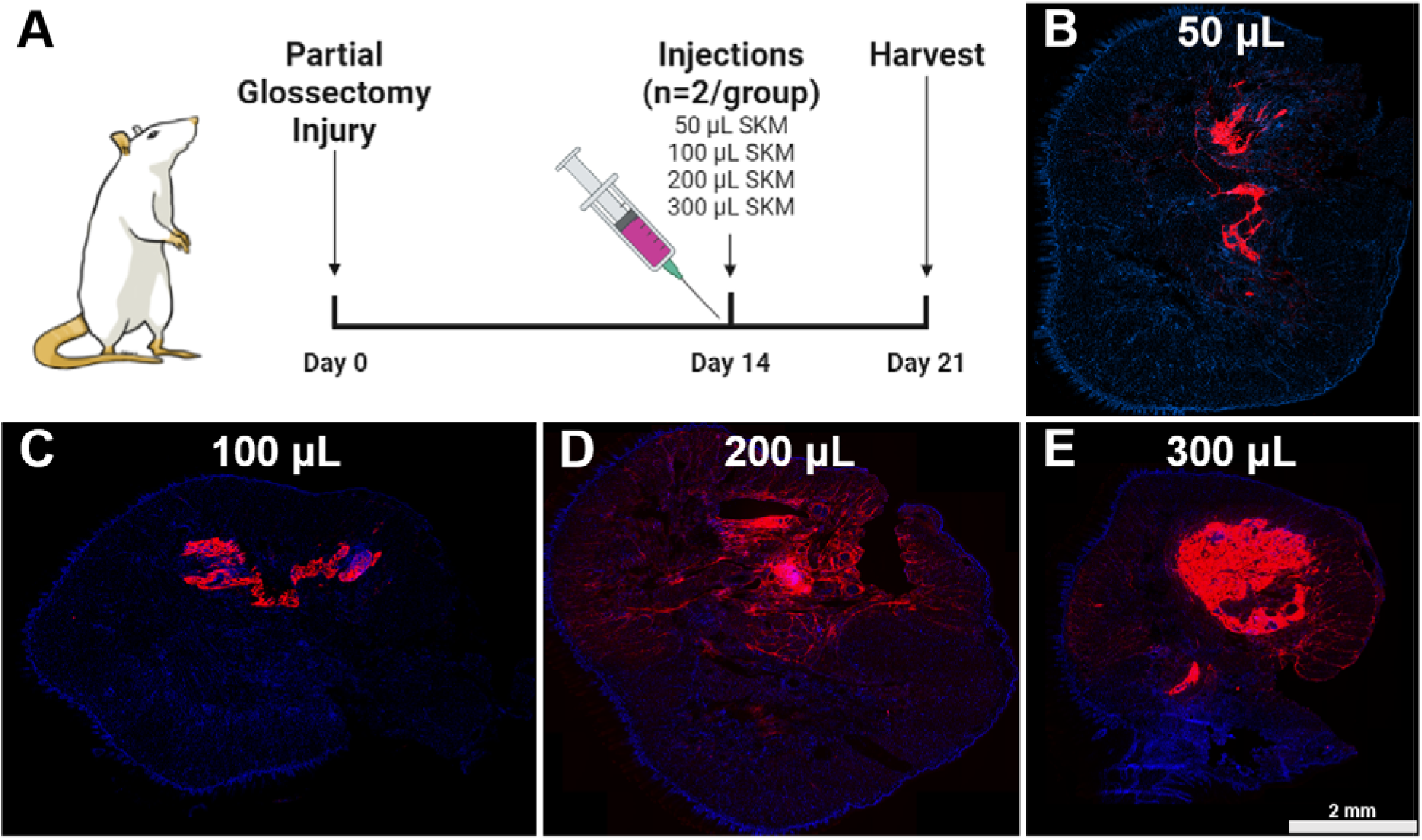
SKM hydrogel injection volume optimization after tongue partial glossectomy injury. (**A**) A range of SKM volumes were injected 2 weeks after partial glossectomy injury, and tissues were harvested 1 week post-injection. Representative fluorescent images show pre-labeled SKM material in red against a DAPI counterstain in blue to indicate cell nuclei. SKM material spread and retention in the target tissue is shown for a range of injection volumes including 50 (**B**), 100 (**C**), 200 (**D**), and 300 μL (**E**). Scale bar: 2 mm.

Based on previous ECM hydrogel studies, greater material retention led to improved repair with a typical degradation time of approximately 3 weeks; thus, it was desirable to see high material retention in the tissue at 1 week post-injection to enable prolonged pro-regenerative cues and sufficient repair.^18,21^ Representative images of injection boluses are shown for all SKM dosages (Figure 1B-E), demonstrating increased material retention and spread through the tongue with higher injection volumes. Therefore, we found that 200 and 300 μL injection volumes were suitable for further investigation.

### Higher dose of SKM hydrogel reduces scar formation in a rat model of tongue fibrosis

To investigate therapeutic potential of SKM hydrogel, the chosen injection volumes – 200 and 300 μL – were injected 2 weeks following partial glossectomy injury (Figure 2A). The experimental control group received a 200 μL saline injection at the same time point. The 300 μL volume was initially anticipated to be inferior to 200 μL because we hypothesized that the larger injection would be disruptive to the tissue. Thus, 200 μL of saline was injected as experimental control. Whole tongues were harvested 4 weeks after injection for assessment of histomorphological properties. First, we investigated whether SKM injection decreased scar formation in this injury model, which is important as fibrosis contributes to dysphagia following surgical resection and radiation of head and neck cancers. Representative brightfield images of tongue cross-sections from the treatment groups are shown in Figure 2B-D. As shown in Figure 2E, the scar area fraction was significantly reduced in the 300 μL SKM group compared to both 200 μL SKM (P=0.005) and 200 μL saline (P=0.02) groups.

**Figure 2.**
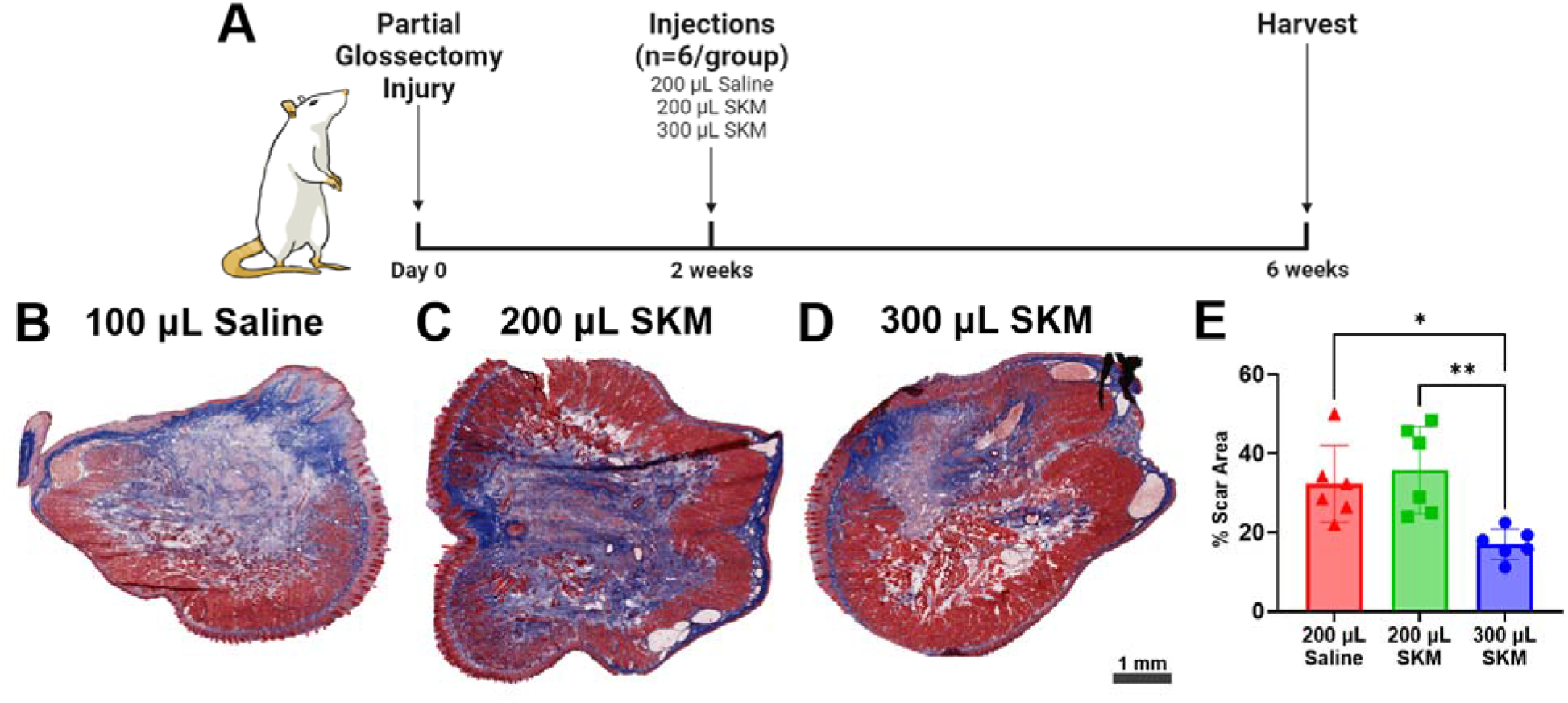
SKM injection after partial glossectomy injury reduces scar formation. (**A**) Timeline of study to assess therapeutic potential of two doses of SKM hydrogel injection. Tissues were harvested 4 weeks after injection for assessment of histomorphological properties. Short-axis tongue cross sections were stained with Masson’s Trichrome, in which collagen is blue, denoting the scar region, and muscle fibers are red. Representative images for the 300 μL SKM (**B**), 200 μL SKM (**C**), and 200 μL saline (**D**) injection treatment groups are shown. Scar area was quantified and normalized to the total cross-sectional area for each section, and values for all sections including the scar were averaged for each animal. (**E**) Treatment groups were compared using one-way ANOVA with Tukey multiple comparisons test (*p<0.05; **p<0.01).

### SKM hydrogel improves muscle regeneration within scar region

For this experiment, the 200 μL SKM group was not assessed, as it did not demonstrate reduction of scar formation; only tissues from the 300 μL SKM and saline groups were used. Short axis cross-sections of tongue specimens were stained with antibodies against collagen I and myofibers membranes (α-sarcoglycan), and a nuclear stain (DAPI). Representative fluorescent images are shown in Figure 3A-B. A higher magnification image demonstrates the presence of skeletal muscle fibers inside the scar area (Figure 3C). To assess the effect of SKM injection on muscle regeneration, we quantified fiber number and cross-sectional area inside the scar region. Although no quantitative differences were observed between the groups (P=0.86), fiber cross-sectional area was significantly greater in the 300 μL SKM compared to saline group (p<0.0001). The percentage of centrally nucleated fibers was quantified to determine if the muscle was actively regenerating; there was no difference in this percentage between SKM and saline treated animals, indicating that muscle regeneration was not ongoing and fiber area measurements were reflective of the tissue at homeostasis.

**Figure 3.**
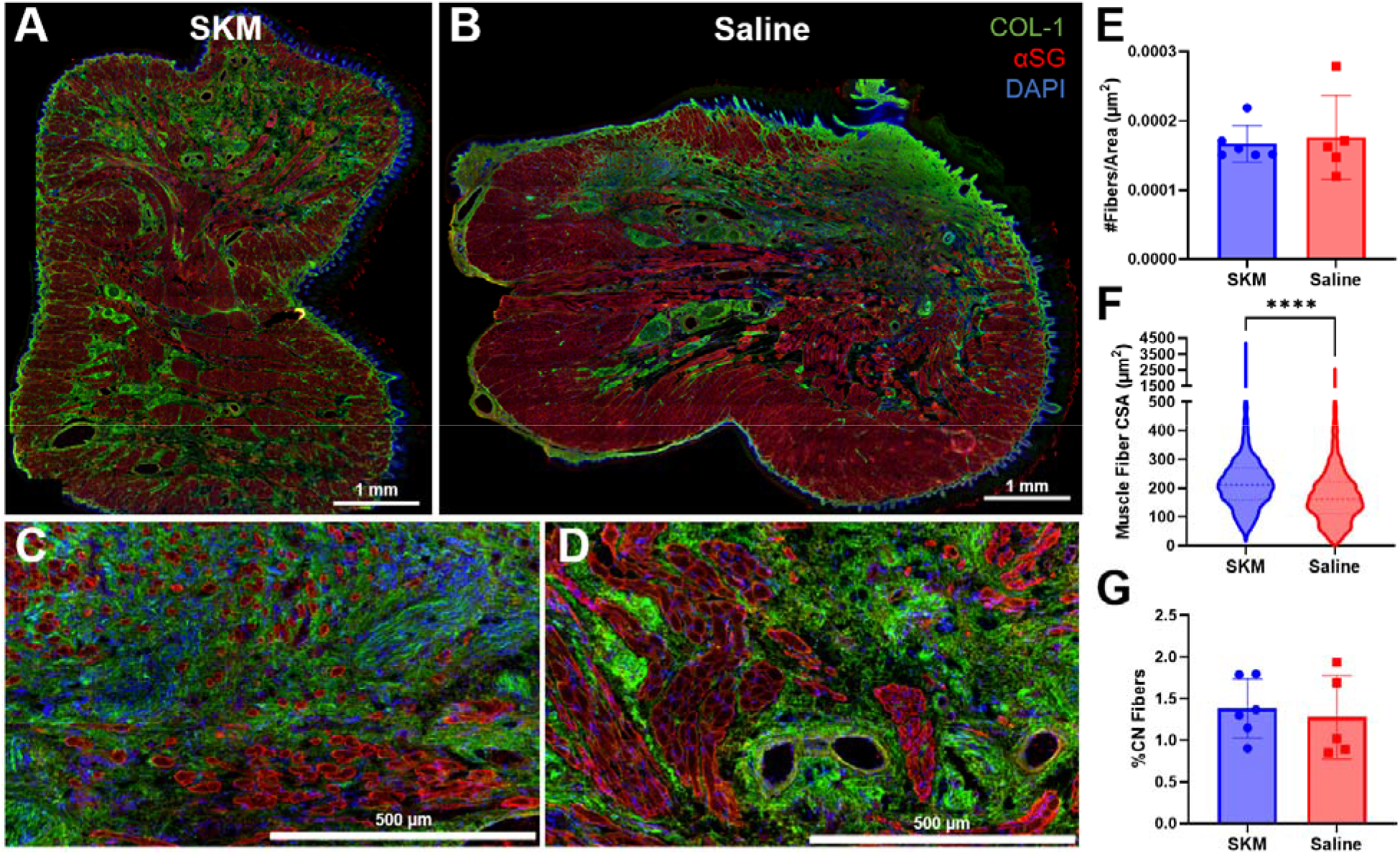
SKM Injection increases muscle fiber area within scar area. Tongue sections underwent immunohistochemical staining to denote the scar (collagen I, green), muscle fiber membranes (α-sarcoglycan, red), and nuclei (DAPI, blue). Representative fluorescent images are shown for 300 μL SKM (**A**) and 200 μL saline (**B**) injection groups (Scale bar: 1 mm). Higher magnification images of (**C**) and (**D**), respectively, demonstrating differences in fiber size and organization within the region of scar. Scale bar: 500 μm. (**E**) Fiber counts were normalized to scar area, showing no significant difference between treatment groups. (**F**) Muscle fiber area distributions are displayed as violin plots, with the dashed line indicating the median. Using a Mann-Whitney t-test, SKM demonstrated significantly increased fiber areas compared to saline injection (****p<0.0001). (**G**) Within the scar area, proportion of fibers with centralized nuclei did not differ between groups.

### SKM hydrogel upregulates pro-regenerative immune response and induces angiogenic signaling

Having demonstrated significant histomorphological improvements with SKM injection in this model, we then conducted a gene expression study to investigate potential mechanisms driving these changes. Two weeks following partial glossectomy injury, 300 μL of SKM or saline was injected into the site of injury. Additionally, non-injected injured controls were used. To capture physiologically relevant timepoints for early changes in the immune response, myogenesis, and ECM remodeling, tissues were harvested for RNA isolation at 3 and 7 days post-injection (Figure 4A). A custom NanoString multiplex gene expression panel of 145 genes for rat skeletal muscle, including pathways of interest such as the immune response, myogenesis, muscle anabolism/catabolism, angiogenesis, and ECM remodeling, was used to characterize the tissue response to SKM injection.

**Figure 4.**
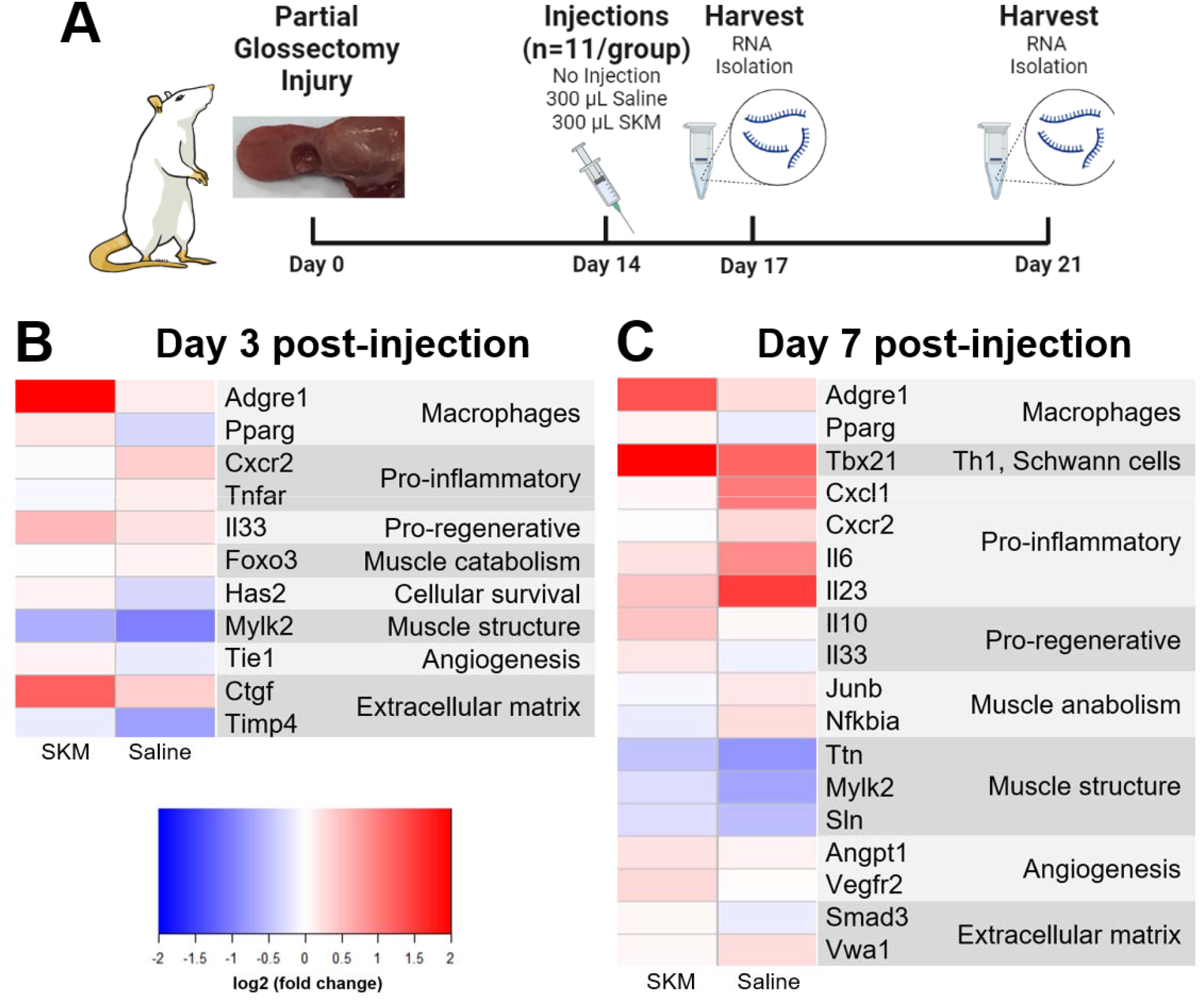
SKM hydrogel injection induces differential regulation of genes involved in the immune response, muscle catabolism and anabolism, muscle structure, angiogenesis, and ECM remodeling. (**A**) Experimental timeline for gene expression study. Tissues were harvested at 3 and 7 days post-injection for RNA isolation, after which a custom rat skeletal muscle panel of 145 genes was analyzed with the NanoString multiplex assay. Gene expression data from SKM and saline controls were normalized to non-injected controls. Heatmaps were generated for genes which were differentially expressed between SKM and saline groups, and data are presented as average fold changes with respect to non-injected controls. Heatmaps for differentially expressed genes at day 3 (**B**) and day 7 (**C**) post-injection are shown.

The significant differentially expressed genes between SKM and saline are listed for days 3 and 7 post-injection alongside their associated pathways in Figures 4B and C, respectively. To visualize the relative differences in expression for each gene between SKM and saline groups, heatmaps were generated using fold changes with respect to non-injected injured controls. At day 3 post-injection, SKM led to upregulation of *Adgre1*, a macrophage marker, and *Pparg*, a macrophage transcriptional marker of pro-repair macrophages.^22^ We further observe SKM immunomodulation through the downregulation of pro-inflammatory chemokine *Ccr2* and upregulation of pro-regenerative cytokine *Il33* with respect to saline. SKM injection upregulated expression of *Has2*, a hyaluronic acid synthase, and *Mylk2*, involved in skeletal muscle contraction, compared to saline injection. SKM also resulted in upregulation of *Ctgf*, which is known for a pro-reparative role in wound healing when expressed transiently in early injury response.^23^ Compared to saline, SKM group had higher expression of *Timp4*, a matrix metalloproteinase inhibitor that regulates ECM remodeling as well as angiogenesis.^24^ *Tie1*, another gene involved in angiogenesis, was also upregulated in SKM with respect to saline.

At day 7 post-injection, SKM upregulation of macrophage markers was sustained (*Adgre1*), and we additionally observed upregulation of *Tbx21*, a marker for type 1 T-helper cells. Chemokines (*Cxcl1, Cxcl2*) and cytokines (*IL6, IL23*) associated with a pro-inflammatory response were downregulated in SKM compared to saline, while cytokines associated with an anti-inflammatory or pro-regenerative response (*IL10, IL33*) were upregulated. While SKM did not upregulate genes related to muscle anabolism (*Junb, Nfkbia*), we observed an upregulation of genes associated to muscle structure (*Ttn, Mylk2, Sln*) in the SKM group as compared to saline. Finally, we observed that SKM induced upregulation of angiogenesis genes *Angpt1* and *Vegfr2*. Overall, the gene expression data from both day 3 and 7 post-injection are consistent in demonstrating that SKM injection reduces pro-inflammatory response, increases pro-regenerative response, and increases angiogenic signaling.

## Discussion

Dysphagia is a life-threatening and morbid sequela following treatment of head and neck cancers. However, clinical approaches to dysphagia are limited primarily to rehabilitative exercises, which lack efficacy and particularly have not been shown to improve tongue strength and swallowing function. Cell therapies under investigation have not demonstrated significant improvement in tongue function clinically and pose significant challenges for translation, given difficulties with cost, manufacturing, and logistics of cell therapies. There is a need for costeffective and easily administered therapeutics that can achieve tissue regeneration to revert the muscle atrophy and fibrosis associated with head and neck cancer treatment in order to improve swallow function.

SKM hydrogels have previously demonstrated induction of muscle regeneration, neovascularization, and ECM remodeling in several skeletal muscle conditions, including ischemia reperfusion^17^ and birth-related mechanical injuries.^19^ Thus, we hypothesized that injection of SKM hydrogel could mitigate scar formation and improve muscle regeneration in an animal model of tongue fibrosis, which was previously developed to model pathologies associated with head and neck cancer treatment.^9^ In this study, we demonstrated that injection of SKM, when delivered two weeks following a partial glossectomy injury, significantly improved histomorphological properties of tongue tissues. The active phase of muscle regeneration was completed 4 weeks after injection, as demonstrated by the proportion of muscle fibers with centralized nuclei in the site of injury,^25^ so this timepoint was used for evaluation of histolomorphological changes in the tongue. At this timepoint, we found that 300 μL SKM injection reduced scar formation and increased muscle fiber area within the scar region. These data suggest therapeutic efficacy of SKM in promoting constructive remodeling and reversing fibrosis of the tongue consequent to surgical resection of the tissue. Interestingly, a smaller injection volume of SKM did not significantly reduce scar area as compared to saline controls. This is likely due to the shorter retention time with the smaller volume, which was insufficient to induce a therapeutic effect and extensive histomorphological changes.

Consistent with previous studies on ECM hydrogels,^19,26^ we observed that SKM injection downregulated pro-inflammatory genes with a concurrent upregulation of chemokines and cytokines associated with a pro-regenerative response. We observed an upregulation of Tbx21, which is commonly used as a marker for Th1 cells, but without concomitant upregulation of Th1 cytokines (IFNγ/, TNFα, IL-2, IL-12). Tbx21 is, however, also expressed in perisynaptic Schwann cells and given the lack of Th1 cytokines, may instead suggest an improvement in reinnervation of the injured muscle^27^. Genes related to other pro-inflammatory cytokines and chemokines were downregulated in SKM, while pro-regenerative or anti-inflammatory cytokines were upregulated. *Il10* and *Il33* are particularly associated with muscle repair, M2 macrophages, T_reg_ cell activity, and potentially pro-myogenic activity of fibro-adipogenic progenitor cells.^28,29^

Additionally, we found that SKM injection upregulated genes involved in angiogenesis, which supports our previous studies that determined SKM induces neovascularization in other skeletal muscle injury models.^17,21^ While we did not observe significant modulation in gene expression related to myogenesis, which was expected based on observed histomorphological changes, SKM did induce upregulation of hyaluronic acid synthesis, which is transiently expressed during hypertrophy,^30^ and other genes related to myofiber contractile proteins. Changes in myogenesis have been observed at 7 days post-injection of SKM in a mechanical birth injury model.^19^ In contrast, ECM scaffold implants have demonstrated limited improvements in myogenesis following VML injury.^31^ Thus, while we had initially thought that SKM injection would induce myogenesis based on other injury models, SKM injection appears to influence tissue repair through alternate mechanisms in VML-type injury. Additionally, we did not observe significant changes in ECM remodeling pathways. Potentially, the early timepoint of transcription analysis, which was intended to capture changes in the immune response may have preceded significant alterations in ECM remodeling, based on previous VML studies.^32^ Further investigation into transcriptional changes in other pathways, such as fibro-adipogenic progenitor^28^ or pericyte^33^ cell activity – populations that may contribute to fibrogenic or myogenic responses – may further characterize SKM’s mechanism of action in this injury model.

One limitation of the partial glossectomy injury model is that while it induces muscle damage and scar formation that mimic the pathologies associated with head and neck cancer treatment, it does not include radiation. Future studies are necessary to include irradiation for a potentially more severe injury model to further evaluate the efficacy of SKM injection. Additionally, the NanoString multiplex gene expression study is a biased approach; we assayed specific sets of genes related to the pathways mentioned, but it would be advantageous to use an unbiased approach that facilitates potential discoveries of new mechanisms of action that govern the pro-regenerative effect of SKM.

Overall, the data support our hypothesis that SKM is a promising therapeutic for the treatment of fibrosis in the tongue. In this rat model of tongue fibrosis, we demonstrate significant histomorphological improvements following SKM injection. Our gene expression data further suggest immunomodulation towards a pro-regenerative phenotype and improvement of angiogenesis, which is important for vascularization of the damaged tissues. This study encourages further investigation of SKM for the regeneration of damaged tongue tissue for the potential treatment of dysphagia.

## Conclusions

In conclusion, our study demonstrates that a cost-effective and easily administered acellular tissue-specific biomaterial significantly reduces scar formation and improves muscle regeneration in a rat model of tongue fibrosis following partial glossectomy. We provide evidence that this biomaterial is immunomodulatory and upregulates genes related to angiogenesis. Overall, this biomaterial, which may be delivered minimally invasively, presents a promising option for future clinical translation in the regeneration of tongue muscle and treatment of dysphagia.

## Notes

### Competing Interest Statement

The authors have declared no competing interest.

