## Supplemental Tables for "Immunomodulatory Extracellular Matrix Hydrogel Mitigates Scar Formation in a Model of Tongue Fibrosis"

**Supplementary**

Table S1: Differentially expressed genes between SKM and Saline at 3 days post-injection

| Gene ID | logFC | pvalue |
| --- | --- | --- |
| Adgr1 | 1.61850664 | 4.8004E-07 |
| Ctgf | 0.70633627 | 0.00414928 |
| Cxcr2 | -0.34528434 | 0.02096618 |
| Foxo3 | -0.12315037 | 0.04937981 |
| Has2 | 0.3513814 | 0.0205203 |
| Il33 | 0.27797367 | 0.00260988 |
| Mylk2 | 0.32294349 | 0.04255447 |
| Pparg | 0.38672879 | 0.00212215 |
| Tie1 | 0.19614778 | 0.00181974 |
| Timp4 | 0.53425493 | 0.00122851 |
| Tnfar | -0.1528032 | 0.02158274 |

Table S2: Differentially expressed genes between SKM and Saline at 7 days post-injection

| Gene ID | logFC | pvalue |
| --- | --- | --- |
| Angpt1 | 0.14253229 | 0.0229001 |
| Adgr1 | 1.12361604 | 4.9195E-10 |
| Il10 | 0.39355048 | 0.01879268 |
| Il23 | -1.05792432 | 0.00867724 |
| Il33 | 0.28718578 | 0.00116824 |
| Il6 | -0.70274986 | 0.04404408 |
| Junb | -0.23130409 | 0.03881962 |
| Mylk2 | 0.43166396 | 0.01019355 |
| Nfkbia | -0.39847947 | 0.00092809 |
| Pparg | 0.24223418 | 0.00903286 |
| Sln | 0.25567115 | 0.04625474 |
| Smad3 | 0.21059108 | 0.00569874 |
| Ttn | 0.35984638 | 0.04332277 |
| Tbx21 | 0.84771311 | 0.01019494 |
| Vegfr2 | 0.30472051 | 0.01561083 |
| Vwa1 | -0.21981367 | 0.04637678 |
| Cxcl1 | -0.98049511 | 0.0487632 |
| Cxcr2 | -0.33182052 | 0.02507791 |
